# Detection of introduced and resident marine species using environmental DNA metabarcoding of sediment and water

**DOI:** 10.1101/440768

**Authors:** Luke E. Holman, Mark de Bruyn, Simon Creer, Gary Carvalho, Julie Robidart, Marc Rius

## Abstract

Environmental DNA (eDNA) surveys are increasingly being used for biodiversity monitoring, principally because they are sensitive and can provide high resolution community composition data. Despite considerable progress in recent years, eDNA studies examining how different environmental sample types can affect species detectability remain rare. Comparisons of environmental samples are especially important for providing best practice guidance on early detection and subsequent mitigation of non-indigenous species. Here we used eDNA metabarcoding of COI (cytochrome c oxidase subunit I) and 18S (nuclear small subunit ribosomal DNA) genes to compare community composition between sediment and water samples in artificial coastal sites across the United Kingdom. We first detected markedly different communities and a consistently greater number of distinct operational taxonomic units in sediment compared to water. We then compared our eDNA datasets with previously published rapid assessment biodiversity surveys and found excellent concordance among the different survey techniques. Finally, our eDNA surveys detected many non-indigenous species, including several newly introduced species, highlighting the utility of eDNA metabarcoding for both early detection and temporal / spatial monitoring of non-indigenous species. We conclude that careful consideration on environmental sample type is needed when conducting eDNA surveys, especially for studies assessing community change.

## Introduction

Anthropogenic activities have widespread impacts on global biodiversity^1,2^ and can negatively affect ecosystem services and function^3^. Cumulatively these actions create an urgent need to develop monitoring tools that rapidly and accurately detect community composition in ecosystems. Existing biodiversity survey techniques have been criticised for their methodological limitations (e.g. observer bias or taxonomic resolution)^4,5^ and are typically standardised by a survey time limit or through reaching asymptote of a species discovery curve^6,7^. Such surveys often focus on the detection of a specific taxonomic group that are being targeted, with no ability to retrospectively separate mis-identified species in light of new species discoveries. This is of critical importance for biodiversity monitoring as an increasing number of studies are revealing the widespread presence of molecular cryptic species (i.e. morphologically similar but genetically distinct species^8^). For example, between 9,000-35,000 marine species (2.7% of the total number of known marine species) are considered molecular cryptic, and genetic studies often reveal widespread marine species containing multiple cryptic lineages^9,10^. This highlights the need to integrate morphological and genetic approaches to accurately detect community composition.

One approach that has the potential to overcome some of the above limitations is the use of nucleic acids found in environmental samples, such as water, soil or sediment, to infer presence or absence of living organisms in the ecosystem^11^. This genetic material, known as environmental DNA (eDNA), is a poly-disperse mixture of tissue, cells, subcellular fragments and extracellular DNA lost to the environment through the normal life and death of organisms^12,13^. Environmental DNA surveys have been used in targeted detection (i.e. single species) studies with qPCR assays^14–17^, and in community (i.e. multi-species) studies using metabarcoding^18–20^. These surveys are highly sensitive and once the methodology is optimised are amenable to automation^21^. However, validity and replicability rely on appropriate experimental design and an understanding of the effects of methodological choices during sampling, sequencing library preparation and bioinformatic analysis^22,23^. Although it is well-established that eDNA surveys are highly informative and can complement other biodiversity monitoring methods^24^, eDNA studies assessing how different sampling techniques affect species detectability remain rare.

One area where accurate monitoring tools are critical is the detection of non-indigenous species (NIS). NIS are those that have been transported through human action from their native range into a novel geographic location. Only a subset of the total number of NIS have a net negative effect^25^ but these pose a severe threat to anthropogenic activities, human health and indigenous biodiversity^26–30^. Most marine NIS have spread globally via vectors such as transoceanic shipping or canals connecting large water bodies^30,31^. At smaller (10s of km) geographical scales, other vectors such as intraregional boating significantly enhance the spread of NIS^32^. In coastal areas, studies have highlighted the importance of monitoring marinas and harbours^6^, as these are hotspots of NIS and together with marine infrastructure (e.g. breakwaters, artificial reefs) promote the spread of NIS^33^. In these habitats, NIS often outcompete native resident species and dominate artificial hard substrata^34,35^. Marinas and harbours have distinct ecological and physico-chemical conditions compared to the surrounding marine environment^36,37^. Consequently, specific sampling and surveying protocols are needed to study marine organisms in these environments, with eDNA surveys offering huge promise for early detection and management of NIS.

Recent work has identified a vast range of protocols for the collection and extraction of eDNA from different environmental sample types (e.g. water, sediment)^38–40^. Despite this progress, we are only just beginning to understand how the choice of environmental sample type affects species detectability^41,42^. For example, we would not expect to detect nektonic in addition to benthic organisms in morphological analyses of a sediment core, but several eDNA studies have detected both of these groups in eDNA isolated from marine sediment^41,43^. Understanding which proportion of the total community is detected using eDNA isolated from different sample types is essential to place eDNA surveys in the context of existing methods, especially when studying NIS.

Here we used eDNA metabarcoding of COI (cytochrome c oxidase subunit I) and 18S (nuclear small subunit ribosomal DNA) genes to compare alpha and beta diversity between sediment and water samples collected in marine urban environments. We then compared the eDNA metabarcoding results with previously published biodiversity data to identify if NIS detection was comparable between methods. We subsequently parsed our eDNA metabarcoding dataset to identify globally relevant NIS and several recently introduced NIS in the study region. We then outlined the strengths and weaknesses of eDNA metabarcoding for the detection of NIS and more broadly community composition. Finally, we discussed how this technique can help conservation efforts for both assessing indigenous biodiversity and mitigating the deleterious effects of NIS.

## Results

### Raw sequencing results and OTU generation

Sequencing produced a total of 17.8 million paired end reads, with 15.2 million sequences remaining after paired end read merging and quality filtering. The average number of sequences per sample after filtering (excluding those from control samples) was 200,185 ± 64,019 (s.d.). Negative control samples contained an average of 811 ± 3,402 (s.d.) sequences. One negative control sample contained ~15,000 sequences that mapped to an operational taxonomic unit (OTU) having 100% identity to a sequence of a terrestrial fungi (Genbank Accession number: FJ804151.1), excluding this OTU from the entire analysis resulted in an average of 51 ± 94 (s.d.) sequences per no-template control sample. Denoising produced 8,069 OTUs for COI and 2,433 for 18S with 6,435 and 1,679 remaining respectively after OTU curation with LULU. Taxonomic annotation identified 622 OTUs from the 18S rRNA dataset and 481 OTUs from the COI dataset. Taxonomic data from World Register of Marine Species could be retrieved for 200 of the annotated COI OTUs and 190 of the 18S OTUs.

### Biodiversity detection

The effects of preservation techniques for water eDNA samples differed between the target amplicons. The 18S rRNA amplicon produced significantly more OTUs (Wilcoxon signed-rank test, p < 0.05) in samples preserved by freezing compared to Longmire’s preservation method, while no significant differences (Wilcoxon signed-rank test, p = 0.55) between preservation treatments were observed for the COI amplicon (see Supplementary Information 2 for details). As a conservative approach all subsequent analyses used sample data from the frozen samples. The minimum number of reads per sample was 137,624 and 117,915 for the COI and 18S datasets, respectively, and so samples were rarefied to these numbers of reads. A consistently greater number of OTUs were detected in the sediment samples compared to the water samples across all sites and both markers as shown in Fig 1b,c. In all cases, unique OTUs were detected in both water and sediment samples, but the mean proportion of unique OTUs across 18S and COI detected in water was lower (49.2%) than in sediment (73.8%). A two-way ANOVA testing the effects of sample type, site and their interaction on the number of OTUs indicated a significant effect of the site-sample type interaction (p < 0.001) for both 18S and COI (see Supplementary Information 3 for full model output). Ordination plots based on the Bray-Curtis dissimilarities (Fig. 1d,e) showed that OTUs found in sediment and water eDNA differed in community structure as much as among sites. Additionally, the PERMANOVA model indicated significant differences (p < 0.001) among sites and eDNA sample types in both the 18S and COI datasets (see Supplementary Information 4 for full model output). Accordingly, eDNA sample type in the PERMANOVA model explained 23.2% and 32.5% of the variation in the 18S and COI data respectively, while site explained 34.2% and 30.5% in the COI and 18S data. At phylum level (Fig. 2), taxonomy did not predict the sample type of detections. However, an exact binomial goodness of fit test showed non-random detection proportions in the Nematoda and Platyhelminthes (p < 0.001 and p = 0.038 respectively, see Supplementary Information 5 for full details), with eDNA detections mostly in sediment in both cases.

**Fig. 1.**
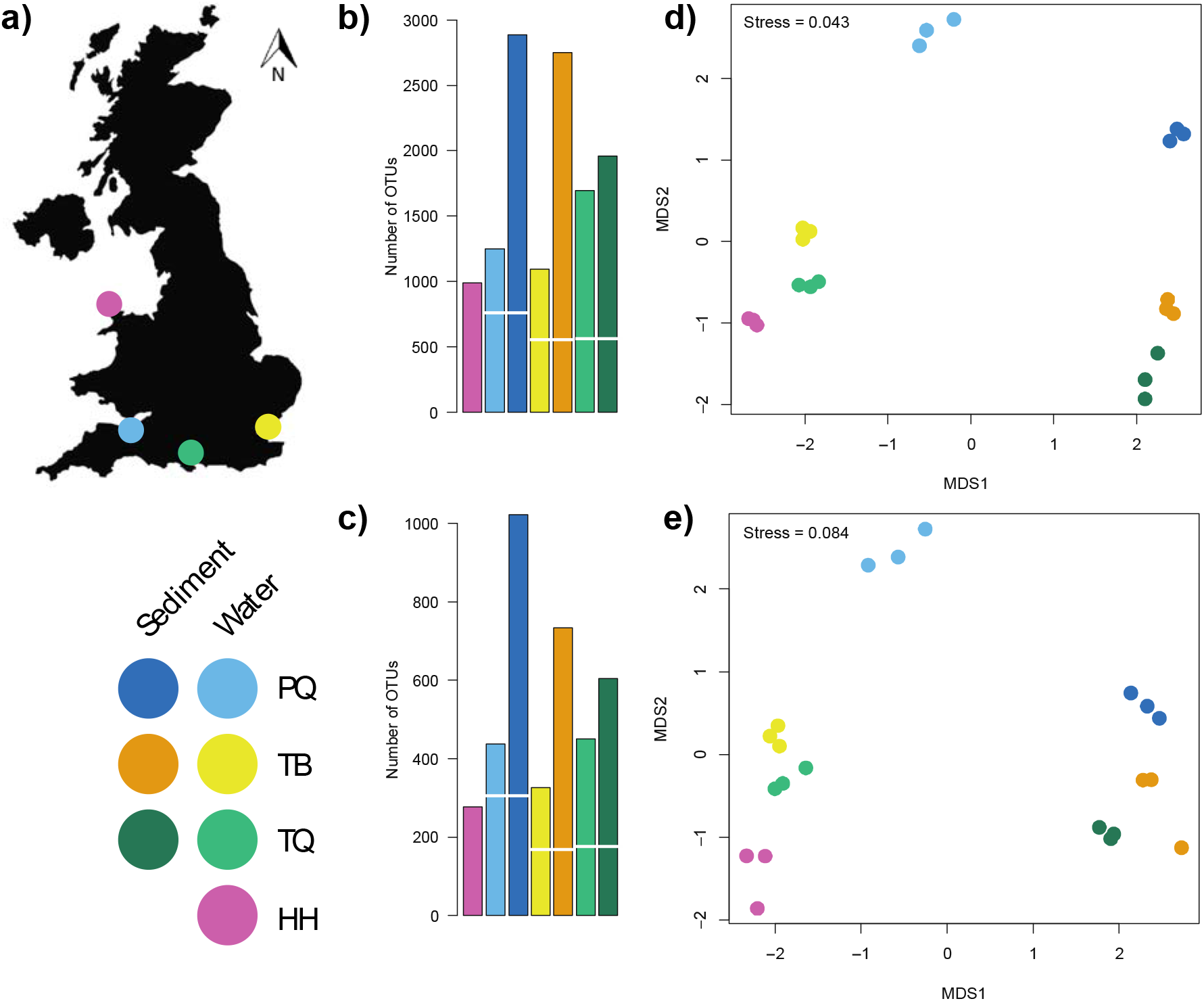
**a** Map of the United Kingdom indicating the geographic position of the sampled sites, a legend indicates the four sites (PQ, TB, TQ and HH) and colours for water and sediment eDNA samples for each site. Barplots detailing number of OTUs (operational taxonomic units) detected across sampling sites and eDNA sample type for **b** COI and **c** 18S rRNA metabarcoding of UK marinas, the break in bars indicates the number of shared OTUs between sediment and water eDNA samples. Non-metric multidimensional scaling ordination plots based on Bray-Curtis dissimilarities between: **c** COI and **d** 18S rRNA metabarcoding of marina sediment and water eDNA samples.

**Fig. 2.**
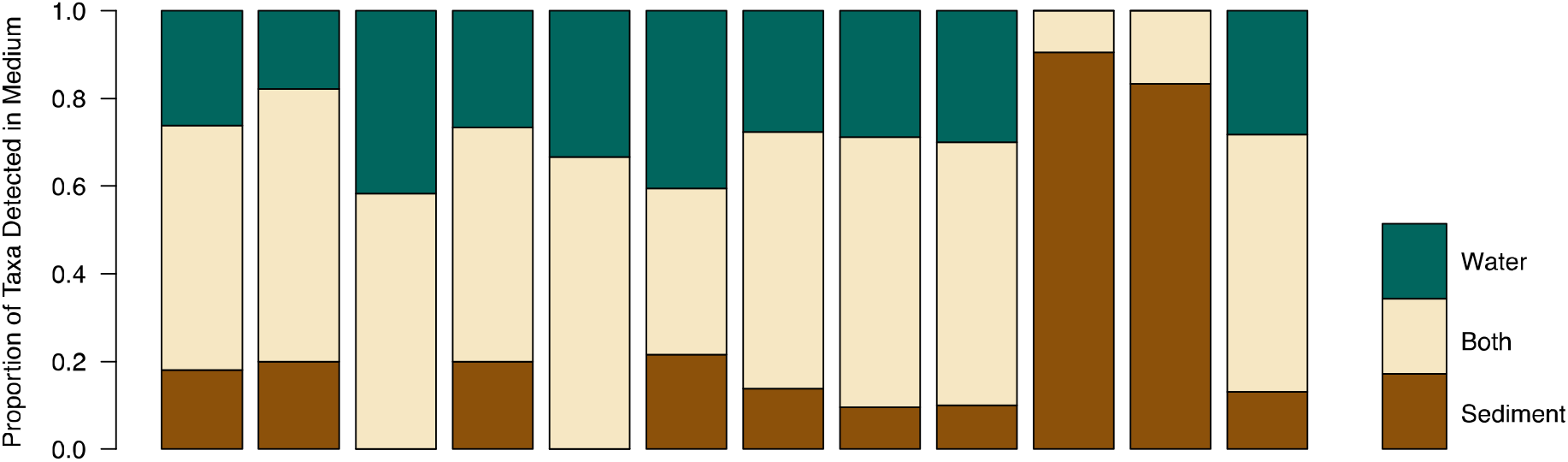
Horizontal stacked barchart detailing proportion of operational taxonomic units detected in eDNA from sediment, water or both sediment and water across the 14 phyla for pooled 18S rRNA and COI metabarcoding of marinas in the United Kingdom.

### Detection of non-indigenous species

As the 18S region lacks the appropriate resolution for taxonomic assignments at species level^44,45^ only the taxonomic assignments from the COI were considered for the identification of NIS. In total 18 NIS to the study region and 24 species documented as NIS in other regions were detected across the four sites (see Supplementary Table 2 for full list). Out of the detected NIS, eight were present in the list of 21 NIS previously detected in rapid assessment (RA) surveys at the sampling sites^46–48^. As shown in Fig. 3, the results of the eDNA surveys closely matched those of the RA surveys. Four detections differed from the RA surveys, a single eDNA detection not seen in RA and three RA detections not seen in eDNA surveys (Fig. 3). Remapping of raw reads from sites with incongruent detections to respective COI regions (Genbank Accessions: *Austrominius modestus* KY607884; *Bugula neritina* KY235450; *Ficopomatus enigmatus* KX840011) found hits for the bryozoan *B. neritina* only (five reads from a single replicate). These reads were lost during data filtering and so did not feature in the final dataset. Three species detections in site TQ represented novel introductions: the detection of *Arcuatula senhousia* (Asian date mussel), the nemertean *Cephalothrix simula* and the oligochaete *Paranais frici.* Targeted visual surveys on tidal mudflats within two kilometres of Marina TQ confirmed the presence of live *A. senhousia* individuals. Furthermore, we generated COI sequences from tissue samples of these individuals (Genbank Accession: MH924820 and MH924821) and these provided full length, high identity matches to both known *A. senhousia* DNA sequences and our eDNA derived OTU sequence (see Supplementary Information 6 for details of DNA barcoding).

**Fig. 3.**
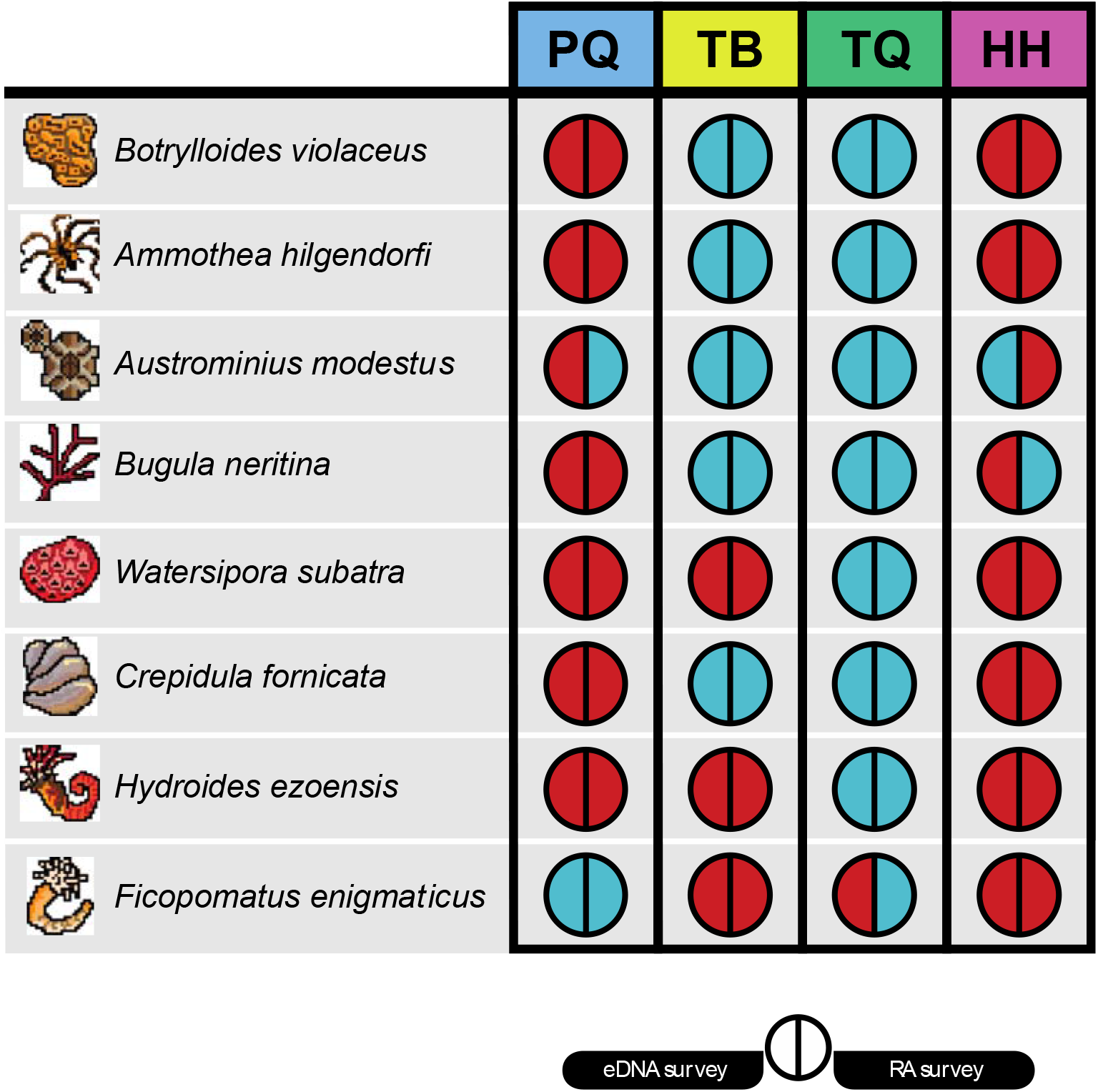
Incidence diagram for seven non-indigenous species across four sampling sites. For each species-location the left semi-circle indicates the detection using eDNA metabarcoding surveys of 18S rRNA and COI fragments, and the right semi-circle indicates the detection from rapid assessment (RA) surveys. Blue indicates a positive detection for that species-location and red indicates no detection.

## Discussion

We demonstrated that the type of environmental sample in eDNA metabarcoding studies affects the measured community composition, indicating that the most comprehensive assessment of biodiversity in a given community comes from the collection of multiple environmental sample types. In addition, we found concordance between our eDNA metabarcoding data and previous biodiversity surveys, demonstrating complementarity of different biodiversity assessment methods. Furthermore, we detected recently introduced NIS, providing support for eDNA metabarcoding as an effective tool for early detection of NIS. This is key as early detection of NIS greatly increases the likelihood of successful control and eventual eradication of NIS. Overall, we demonstrate that type of environmental sample can affect the detection of both whole community composition and particular species of concern.

Our study showed that taxonomic assignments at the level of Phylum did not predict if a species was detected in water, sediment or both environmental sample types (except in Nematodes and Platyhelminthes, whose members are predominantly benthic inhabitants). However, all sampled sites showed higher OTU richness in sediment compared to water. The magnitude of this difference was not fixed across sites, with a significant interaction term in our two-way ANOVA (Supplementary Information 3) indicating that the detected OTU richness differences between sediment and water vary spatially. The majority of research using eDNA to detect aquatic macrofauna has focused on the collection of water samples, while sediment samples have received comparatively less attention (see Figure S1 from Koziol, et al. ^41^). This is surprising considering that sediment samples typically contain three orders of magnitude more eDNA than water^49^. Despite our observations that sediment provided a greater number of OTU richness than water samples, we do not advocate for a particular sample type, as this decision should be driven by the target organisms for a given study. For example, a researcher hoping to use eDNA metabarcoding to measure Nematode diversity, based on our results, should sample marine sediment. Regarding NIS, both water and sediment served as excellent sample types for NIS detection. Consequently, our results suggest that no specific sample type offers a better detection of NIS, likely because NIS are not found in a single taxonomic group. We argue that at a lower taxonomic level the species-specific ecology of eDNA (*sensu* Barnes and Turner ^50^) may lead to convergent eDNA occupancy in different environmental sample types. Further work is needed to clarify how eDNA partitions into adjacent environmental samples across the tree of life. A key unknown is the underlying explanation for eDNA metabarcoding data from sediment samples generating more OTUs in comparison to water. One hypothesis is that eDNA from sediment includes extracellular ‘free’ DNA that is not retained by the filters used to process water for eDNA samples. Studies focussing on eDNA surveys have found little evidence identifying what proportion of total eDNA is extracellular DNA. However, using qPCR Turner, et al. ^12^ identified that eDNA particles with a size of less than 0.2μm, well above the size of extracellular DNA, are less than 10% of the total eDNA pool for a teleost fish species. If this pattern is observed in other metazoans then extracellular eDNA may have little effect on the differences of OTU richness detected here. An alternative hypothesis is that due to the settlement and persistence dynamics of sediment, it contains a greater diversity of eDNA fragments (both extra and intracellular). However, we know very little about these processes and their comparative effects in sediment are unknown.

Current eDNA metabarcoding research has identified large variation in the detected marine biodiversity across small spatial scales (100s of metres) in both sediment^51^ and water^52,53^. Additionally, fractionation of environmental samples (i.e. sorting samples by particle size class) can produce significant differences in the metabarcoding results between fractions^54,55^ indicating significant variation can be found within sites. We found similar patterns, with PERMANOVA modelling indicating that site and environmental sample type contain approximately equivalent variation in OTU dissimilarity. Future research should explore how different sample types and eDNA extraction methods affect the detection of biodiversity, especially as eDNA metabarcoding moves from an experimental technique to a routine monitoring tool^56,57^.

A key gap in our understanding is the rate at which eDNA degrades in sediment and how this affects our observations. In lake sediments, eDNA can be preserved for thousands of years^58,59^, with eDNA being preserved along with deposited sediments so each core represents a timeline through which past biological community communities can be examined^60^. Here we chose to process only the uppermost section of the sampled cores, with the aim of profiling contemporary species composition. Studies are needed to advance our understanding of how eDNA deposits and degrades in marine sediments in order to temporally contextualise sediment samples.

We found that eDNA metabarcoding accurately detected many NIS, as seen in previous studies^18–20^. By comparing our eDNA data to those collected using existing methods we found close congruence in NIS incidence. The false-negative eDNA detection of *B. neritina* was found to be a result of setting specific bioinformatic parameters, showing that choices made during sequence processing can have a significant effect on the detectability of species in eDNA samples. Indeed, this has previously been shown in metabarcoding of bulk tissue samples^61^ and work is urgently needed to determine the effects of bioinformatic parameters, variable primer binding sites and the choice of reference databases on the detection of NIS from eDNA samples. The remaining incongruent detections may be a result of community turnover among the survey dates or phenological changes affecting species distributions. Indeed, marine coastal communities have been shown to shift in community composition across seasons and reproductive cycles^62,63^. Therefore, our data suggest that in order to enhance existing monitoring programmes, replicated eDNA metabarcoding surveys over time should be performed. In this study we identified several recently introduced NIS in the United Kingdom and confirmed the eDNA detection with targeted local surveys for one NIS. The case of *A. senhousia* is particularly relevant as it is spreading globally^64^ and has the potential to dramatically alter benthic biodiversity when invasive^65,66^. This species produces a cocoon of byssus thread that at high densities (>1,500 individuals/m^2^) interlinks between individuals to form a continuous byssal mat which displaces local eelgrass and native bivalves^67,68^. Recent field surveys along the south coast of the United Kingdom have independently confirmed the presence of both *A. senhousia*^*69*^ and *C. simula*^*70*^. These results confirm the accuracy of eDNA surveys presented here and highlight the benefits of implementing molecular technologies for routine monitoring programmes. As the cost of sequencing continues to decrease and methods improve across the metabarcoding workflow^71^ natural resource managers and researchers will have access to much greater resolution data at a fraction of the cost and time of current monitoring surveys. However, NIS can be missed in surveys based solely on eDNA (e.g. Wood, et al. ^72^) and eDNA studies can detect rare species that are often missed using other methods^73^. Detection of NIS could be further facilitated through autonomous sampling and eDNA surveys^21^ to provide live species incidence data in introduction hotspots, such as ports or marinas. Additionally combining these techniques with eDNA biobanking^74^ could provide an eDNA reference database for specific geographical regions of high biosecurity risk, providing an invaluable resource for both biodiversity managers and researchers to examine the process of biological invasion through time. Taken together, our study shows eDNA metabarcoding to be an effective tool for the detection and identification of both native and NIS from different environmental samples.

## Methods

### Study sites

Four marinas were selected from around the United Kingdom (Fig. 1a) to represent variation in modelled invasion potential^75^, presence of NIS^76^ and benthic habitat type^77^. All chosen marinas have been surveyed previously, so there is a good understanding of the species found in these sites^46–48^. Marina access was contingent on anonymity and so marina names and exact locations are not provided, with Fig. 1a showing approximate locations only. Marina *TQ* is an open marina subject to tides and varying salinity, marina *PQ* is a loch marina open during high tide, and marinas *TB* and *HH* are permanently open to the North and Celtic Sea respectively.

### Environmental DNA sampling

Surveys were conducted during May 2017 (see Supplementary Table 1 for site details) and 24 sampling points were randomly selected within each site. At each sampling point 50ml of water was collected from 10cm below the surface using a sterile 60ml Luer lock syringe and filtered through a 0.22μm polyethersulfone Sterivex filter (Merck Millipore, Massachusetts USA). After collecting seawater from eight sampling points (400ml total volume) the filter was changed, resulting in a total of three filters per site. Pooling of water samples was performed to provide three filter replicates per site that represented the heterogeneity of eDNA in the marina. In order to test the effect of different sample preservation methods, water samples were collected in duplicate in each sampling point. One set of three filters had ~1.5ml sterile Longmire’s solution (100mM Tris,10mM EDTA, 10mM NaCl, 0.5% SDS) applied in the inlet valve^78^. The second set of three filters was kept on ice for no longer than eight hours before being frozen at −20°C. In addition to the water samples, a subtidal sediment sample was collected at the first water sampling point and then after every three water samples, accounting for a total of nine sediment samples per site. We used a UWITEC Corer (UWITEC, Mondsee, Austria) to collect a sediment core of 600mm high and 60mm diameter. We then used a sterile disposable spatula to collect a subsample of 10-20g of sediment from the top 2cm of the core, avoiding sediment collection from the sides of the core. The subsamples were stored in sterile plastic bags and kept on ice for no longer than eight hours before being frozen at −80°C. Due to a malfunction of the corer, no sediment sample was collected in Site HH. Disposable gloves were changed after collection of each sample. All reused equipment was soaked in 10% bleach and rinsed in DNase-free sterile water between sites.

### eDNA extraction

DNA extractions were performed in a PCR-free clean room, separate from main laboratory facilities. No high copy templates, cultures or amplicons were permitted in this clean laboratory. DNA extractions from water samples followed the SX_CAPSULE_ method by Spens, et al. ^39^. Briefly, preservative solution was removed from the outlet and filters were dried at room temperature for two hours. We then added 720μl Qiagen buffer ATL (Qiagen, Hilden, Germany) and 80μl Proteinase K to the filter and all samples were digested overnight at 56°C. After digestion, samples were processed using the Qiagen DNeasy Blood and Tissue Kit as per manufacturer instructions, with a final elution of 200μl PCR grade water.

Sediment extractions were conducted using the Qiagen DNeasy Powermax Soil Kit following the manufacturer’s protocol. The nine samples collected in each site were randomly mixed to form three pooled samples; 10g of pooled sample was processed for the extraction. A total of ten samples were processed, three from each site with a single extraction control.

### Inhibition testing

To ensure extracted DNA was free of PCR inhibitors, a Primer Design Real-Time PCR Internal Control Kit (PrimerDesign, Southampton, United Kingdom) was used. We performed qPCR reactions on each extracted DNA sample following the manufacturer’s protocol with 12.5μl reaction volumes. A positive detection of inhibition due to co-purified compounds from DNA extraction protocols would produce an increase in cycle threshold number in comparison to no template controls. All samples were successfully processed and no samples showed indication of PCR inhibition.

### Primer selection and library preparation

Two sets of primers were chosen for metabarcoding the environmental samples: a 313bp section of the standard DNA barcoding region of the cytochrome c oxidase subunit I gene (COI) using primers described in Leray, et al. ^79^; and a variable length target of the hypervariable V4 region of the nuclear small subunit ribosomal DNA (18S) using primers from Zhan, et al. ^80^. These two primer sets allow for broad characterisation of marine metazoan diversity. Sequencing libraries were prepared using a 2-step PCR approach as detailed in Bista, et al. ^81^. Briefly, this method first amplifies the target region in PCR 1 annealing universal adapters, and then sample specific indices and sequencing primers are annealed in PCR 2. In contrast to Bista, et al. ^81^ we used unique dual-matched indexes for PCR 2 to avoid index crosstalk associated with combinatorial indexing^82^. PCR 1 was prepared in a PCR–free room separate from main laboratory facilities. PCR 1 reactions were conducted in 20μl volumes containing 10μl Amplitaq GOLD 360 2X Mastermix (Applied Biosystems, California, USA), 0.8μl (5 nmol ml^−1^) of each forward and reverse primer and 2μl of undiluted environmental DNA extract. The reaction conditions for PCR were an initial denaturation step at 95°C for 10 minutes followed by 20 cycles of 95°C for 0:30, variable annealing temp (46°C for COI and 50°C for 18S) for 0:30, and extension at 72°C for 1:00. A final extension at 72°C was performed for 10 minutes. The PCR product was cleaned using AMPure XP beads (Beckman Coulter, California, USA) at a 0.8 beads:sample ratio following manufacturer’s instructions. PCR 2 reactions were conducted in 20μl volumes containing 10μl Amplitaq GOLD 360 2X Mastermix, 0.5μl (10 nmol ml^−1^) of both forward and reverse primers and 5μl of undiluted cleaned PCR1 product. PCR conditions were an initial denaturation step at 95°C for 10 minutes followed by 15 cycles of 95°C for 0:30, annealing at 55°C for 0:30, and extension at 72°C for 1:00. A final extension at 72°C was performed for 10 minutes. PCR 2 products were cleaned using AMpure XP beads as above and normalised according to their fluorescence using the Qubit HS Assay Kit (Thermofisher Scientific, Massachusetts, USA). These normalised samples were pooled at an equimolar concentration and then quantified as per manufacturer’s instructions using the NEBNext Library Quant qPCR kit (New England Biolabs, Massachusetts, USA).

Blank filters, DNA extraction kits and positive controls were collected, extracted and sequenced identically to non-control samples (detailed in Supplementary Information 1). Negative controls cannot be meaningfully normalized and thus they were added to the pooled libraries without dilution. The final library was sequenced using an Illumina MiSeq instrument (Illumina, San Diego, USA) with a V3 2 × 300bp kit.

### Bioinformatic analyses

Samples were demultiplexed using the Illumina MiSeq control software (v.2.6.2.1). The demultiplexed data was analysed using a custom pipeline written in the R programming language^83^ (hosted at https://github.com/leholman/metabarTOAD). The steps are as follows. Forward and reverse paired end reads were merged using the -fastq_mergepairs option of USEARCH v.10.0.240^84^ with maximum difference of 15, percent identity of 80% and quality filter set at maximum expected errors of 1. Both the forward and reverse primer sequences were matched using Cutadapt v.1.16^85^ and only sequences containing both primer regions were retained. Sequences were discarded if they were outside of a defined length boundary (303-323bp for COI, 375-450bp for 18S) using Cutadapt. Sequences were then pooled, singletons were discarded and sequences were quality filtered with a maximum expected error of 1 using the -fastq_filter option of vsearch v.2.4.3^86^. Sequences were then denoised and chimeras filtered using the unoise3 algorithm implemented in USEARCH. The resultant operational taxonomic units (OTUs) were curated using the LULU package v.0.1.0^87^. An OTU by sample table was produced by mapping the merged and trimmed reads against the curated OTUs using USEARCH, with the raw query read assigned to the OTU with the best match (highest bit score) within 97% identity. The OTU × sample table was filtered in R as follows. To minimise the chance of spurious OTUs being included in the final dataset any record with less than 3 raw reads were changed to zero and any OTU that did not appear in more than one sample was removed from the analysis. OTUs found in negative controls were removed from the analysis.

### Taxonomic assignment

Assigning correct taxonomy to an unknown set of DNA sequences can be challenging as reference databases are incomplete, contain errors and the taxonomy of some marine groups is uncertain. With such limitations in mind, we assigned taxonomy using a BLAST v.2.6.0+ search^88^ returning the single best hit (largest bit score) from databases within 97% of the query using a custom R script to parse the raw blast results. In the case of multiple sequences attaining equal bit scores for a given OTU an assignment was only made if all reference sequences belonged to the same species. The MIDORI database (UNIQUE_20180221)^89^ was used for the COI data and the SILVA database (SSU r132, subset to contain only Eukaryotes)^90^ was used for the 18S rRNA data. The match taxa tool from the World Register of Marine Species^91^ was used to filter the data to include only marine species and check the taxonomic classification. The World Register of Introduced Marine Species^92^ contains a range of peer-reviewed and technical reports on the global introduced status of a large number of species, we used the online match taxa tool to determine the non-indigenous status of annotations that could be assigned taxonomy from the World Register of Marine Species.

### Statistical analyses

All statistical analyses were conducted in R v.3.5.0. The Vegan R package v.2.5.2^93^ was used to rarefy samples to the minimum sample read depth for each amplicon. The number of OTUs per site/condition was calculated as the number of OTUs with a non-zero number of normalized reads after summing the reads across all three site level replicates. To test if there was a significant difference between the number of OTUs generated by sediment and water eDNA, individual non-summed replicate sample data was used to build a two-way ANOVA model with the formula *number_of_OTUs~sedimentorwater*site* implemented in R using the function aov(). Non-metric multidimensional scaling ordination plots were generated from Bray-Curtis dissimilarity values derived using Vegan. A Permutation Analysis of Variance (PERMANOVA)^94^ was performed using the Bray Curtis dissimilarity following the model *dissimilarity_matrix~sedimentorwater*site* implemented in R using the function *adonis* from the *vegan* package. OTUs with taxonomic assignment were separated into those found in sediment, water or both media and the OTUs were then collapsed at the Phylum level to explore taxonomic patterns of detection in water or sediment. Phyla with less than eight OTUs were combined and represented under category named “other”. To test for non-random counts of species detection between water and sediment within taxa an exact binomial test was performed between counts of species detected in water and sediment. The number of species detected in both water and sediment were halved and the value added to the counts for each sample type with non-integer values conservatively rounded down to the nearest whole number. A correction for multiple comparisons^95^ was applied across the p values from the exact binomial tests generated by the R function binom.test(). Records from rapid assessment surveys previously conducted for non-native invertebrates at the sample sites^46–48^ were compared with the detected species from metabarcoding data.

## Supporting information

Supplementary Table 1 – Site Metadata

Supplementary Table 2 – Non-Indigenous Species

Supplementary Table 3 – Sequencing Library Metadata

## Acknowledgements

We are grateful to John Bishop and Chris Wood from the Marine Biological Association of the United Kingdom for sharing information on marinas and their excellent NIS survey data. We thank the staff of the Environmental Sequencing Facility from the National Oceanography Centre Southampton for advice and assistance during library preparation. We thank Dr Ivan Haigh for assistance in accessing remote sensing data. We acknowledge the Department of Geography and Environment from the University of Southampton for access to coring equipment and laboratory space. LH was supported by the Natural Environmental Research Council (grant number NE/L002531/1).

## Author Contributions

L.E.H. and M.R. designed the experiment, L.E.H. collected samples, generated and analysed the data, designed all figures and wrote the first draft of the paper. L.E.H, M.B., S.C., G.C., J.R. and M.R. contributed critically to further manuscript drafts and gave final approval for publication.

## Data Availability

Raw Illumina sequencing data is available from the European Nucleotide Archive under accession number $$$ (data will be uploaded upon acceptance, available to reviewers upon request).

Associated metadata, script and intermediate files are available online via Zenodo with the following DOI: 10.5281/zenodo.1453958.

