## Supplementary Information 1-6 for "Detection of introduced and resident marine species using environmental DNA metabarcoding of sediment and water"

Mr. Luke E. Holman

Dr. Mark de Bruyn

Prof. Simon Creer

Prof. Gary Carvalho

Dr. Julie Robidart

Dr. Marc Rius

***Supplementary Information* 1. Control Samples**
The following control samples were used. Two sealed filter controls were taken during all field sampling and subject to identical treatment to samples taken during each sampling trip, one contained Longmire’s solution, the second was kept cool during sampling. A filter was opened during DNA extraction to act as an equipment blank for all filters. Distilled water (400ml) was left open in the post-PCR lab for 3 weeks then filtered, this control checked for contaminating amplicons in aerosols in the lab. A blank extraction was used for the DNeasy kit following the same protocol but with no template added. PCR no-template controls were run during PCR1 and PCR2. A positive control sample of 200ml was filtered from a marine tropical aquarium in the reception of the National Oceanography Centre, Southampton, United Kingdom (50.891380N,1.3939209W). The tank contained a variety of hard and soft corals, molluscs and tropical fish. A 400ml sample of seawater was filtered adjacent to the National Oceanography Centre Southampton and 5μl of 1:100 diluted extracted genomic DNA from a species not currently known to United Kingdom waters; *Microcosmus squamiger* Michaelsen, 1927 (Class Ascidiacea, Phylum Chordata) was added to the filter before DNA extraction to act as an inhibition control. This control was used to test if target DNA could be detected after DNA extraction in the presence of inhibitors found in marinas and harbours.

***Supplementary Information 2* Filter preservation treatment**

Raw data was rarefied by the lowest number of reads per sample, this was 117,915 in the 18S dataset and 52,740 in the COI dataset. Detection of an OTU was positive if a non-zero number of normalised reads mapped to an OTU. As shown in Figure S1 below the average number of OTUs detected was always higher in the frozen sample compared to the Longmire’s preserved sample in the 18S data, there was no consistent difference between conditions in the COI dataset. The data was tested for statistical significance using a Wilcoxon signed-rank test. This test showed more OTUs are detected in water samples preserved by freezing in comparison to Longmire’s solution using a 18S amplicon (V=10, p=0.025). No significant difference was found in OTU detection between sample preservation methods using a COI amplicon (V=38, p=0.969).

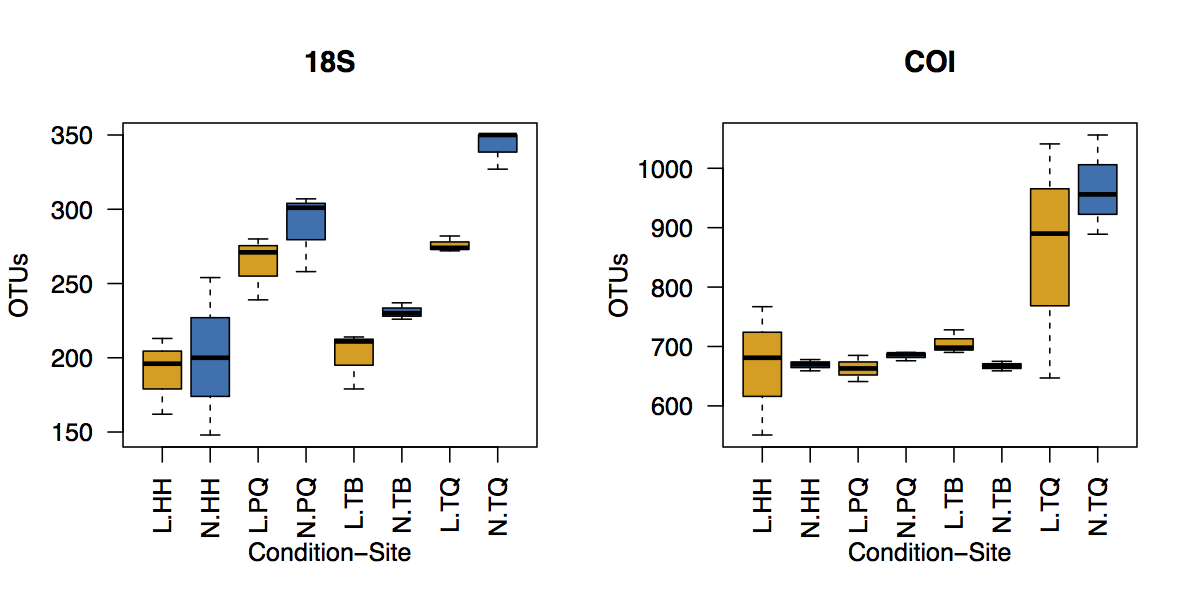

***Figure S1****. Boxplots detailing number of OTUs generated using eDNA metabarcoding of seawater in marina sites with samples preserved with either Longmire’s solution (Yellow) or by Freezing (Blue).*

In order to examine if the effect between preservation methods was driven by low abundance OTUs the analysis was rerun with OTU detection being positive at an increasing threshold of normalised reads from 1 to 200. The results of the Wilcoxon signed-rank test for a significant difference between Longmire’s and frozen OTU detection are shown in Figure S2 below. The results indicated that the significant effect was driven by low abundance OTUs in the 18S dataset. After applying the Bonferroni correction for multiple comparisons, no significant differences remained among conditions.

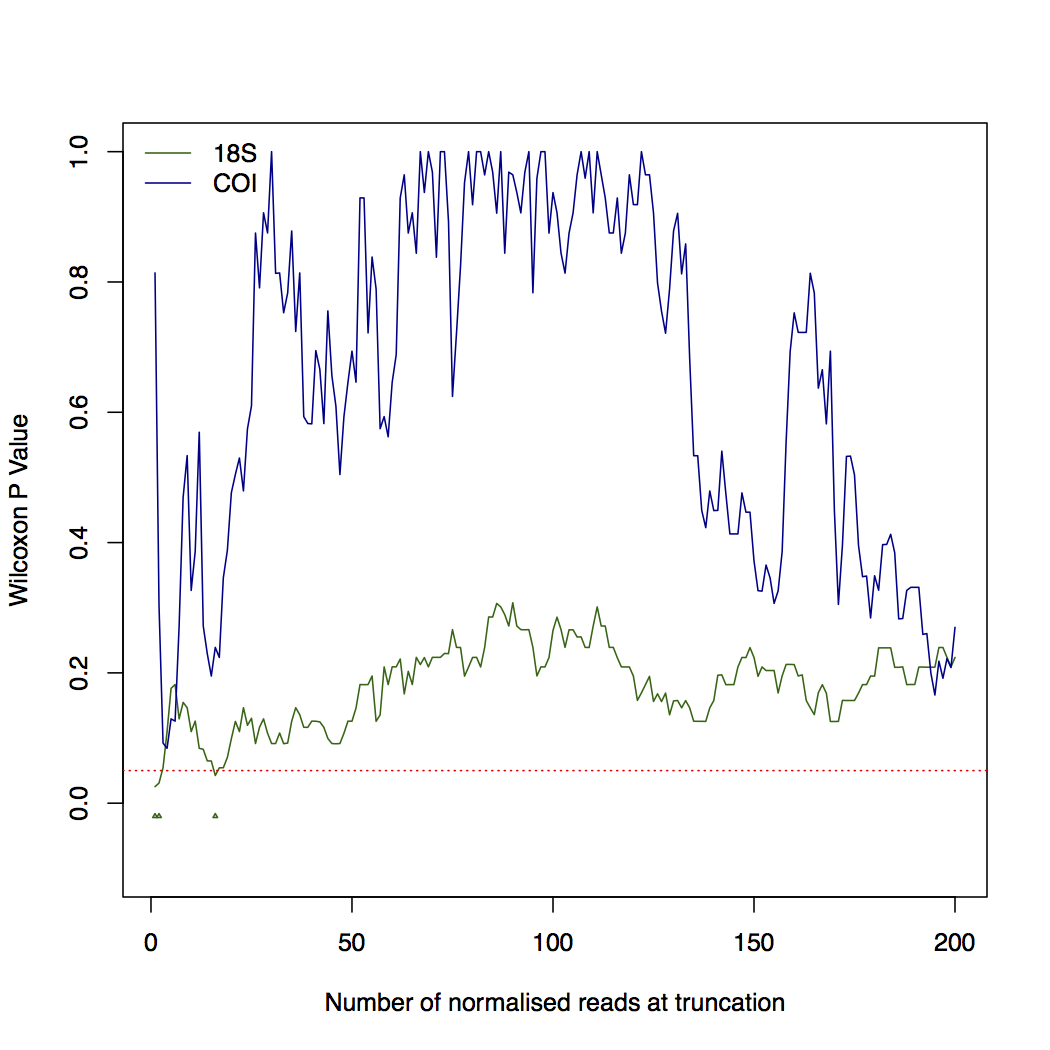

***Figure S2****. Line chart detailing Wilcoxon signed-rank test P value for a difference between the number of OTUs detected across pairs of samples preserved using Longmire’ solution or by freezing as a function of number of normalised reads at truncation. The coloured solid lines indicate the values for a 18S and COI amplicon. The dashed red line marks a value of 0.05 and the coloured points indicate significance at the P<0.05 level.*

Taken together, these results indicated that there is a small but significant difference in the number of OTUs generated in an eDNA metabarcoding experiment between Longmire's and temperature preservation only with the 18S amplicon. This difference is driven by low abundance OTUs which may represent unfiltered false-positive detections or rare sequences.

***Supplementary Information 3*: Model output for linear model with formula *number_of_OTUs~sedimentorwater*site for both 18S and COI metabarcoding of the sampled marinas.***

| **COI** |  |  |  |  |  |
| --- | --- | --- | --- | --- | --- |
|  | Df | Sum Sq | Mean Sq | F Value | Pr(>F) |
| cond | 1 | 4285616 | 4285616 | 171.254 | 1.83E-08 |
| site | 2 | 229337 | 114668 | 4.582 | 0.0332 |
| cond:site | 2 | 1322855 | 661428 | 26.431 | 4.01E-05 |
| Residuals | 12 | 300299 | 25025 |  |  |
| **18S** |  |  |  |  |  |
|  | Df | Sum Sq | Mean Sq | F Value | Pr(>F) |
| cond | 1 | 393089 | 393089 | 98.29 | 3.93E-07 |
| site | 2 | 89435 | 44717 | 11.18 | 0.001813 |
| Cond:site | 2 | 118605 | 59303 | 14.83 | 0.000571 |
| Residuals | 12 | 47989 | 3999 |  |  |

***Supplementary Information 4*: Model output for PERMANOVA model with formula *dissimilarity_matrix~sedimentorwater*site for both 18S and COI metabarcoding of the sampled marinas.***

| **COI** |  |  |  |  |  |  |
| --- | --- | --- | --- | --- | --- | --- |
|  | Df | SumsOfS | MeanSqs | F.Model | R2 | Pr(>F) |
| treatment | 1 | 2.0771 | 2.07708 | 33.96 | 3.25E-01 | 0.001 |
| sites | 2 | 1.9477 | 0.97385 | 15.922 | 0.30508 | 0.001 |
| treatment:sites | 2 | 1.6254 | 0.81271 | 13.288 | 0.2546 | 0.001 |
| Residuals | 12 | 0.734 | 0.06116 | 0.11496 |  |  |
| Total | 17 | 6.3842 | 1 |  |  |  |
| **18S** |  |  |  |  |  |  |
|  | Df | SumsOfSqs | MeanSqs | F.Model | R2 | Pr(>F) |
| treatment | 1 | 1.5626 | 1.5626 | 16.6369 | 0.23213 | 0.001 |
| sites | 2 | 2.2988 | 1.14941 | 12.2377 | 0.3415 | 0.001 |
| Treatment:sites | 2 | 1.7431 | 0.87155 | 9.2793 | 0.25894 | 0.001 |
| Residuals | 12 | 1.1271 | 0.09392 | 0.16743 |  |  |
| Total | 17 | 6.7316 | 1 |  |  |  |

***Supplementary Information 5*: Output of exact binomial test for non-random distribution of number of species detected in sediment or water, number of species detected in water and sediment are reported as ‘N.water’ and ‘N.sediment’ respectively. The raw and multiple-comparison adjusted P values are reported along with the lower and upper 95% confidence intervals.**

|  | N.sediment | N.water | P value | P value adj | Lwr 95% CI | Upr 95% CI |
| --- | --- | --- | --- | --- | --- | --- |
| Platyhelminthes | 11 | 1 | 6.35E-03 | 3.81E-02 | 0.615 | 0.998 |
| Nematoda | 20 | 1 | 2.10E-05 | 2.52E-04 | 0.762 | 0.999 |

***Supplementary Information 6*: Asian Date Mussel Barcoding**

Tissue was sampled and processed in duplicate from the mantle from two *A. senhousia* individuals and immediately subject to DNA extraction using the Qiagen DNeasy Blood and Tissue Kit as manufacturer’s instructions. Each sample was amplified using a set of primers targeting the COI gene (Folmer et al., 1994) conducted in 20μl volumes containing 10μl Amplitaq GOLD 360 2X Mastermix, 0.8μl (5 nmol ml^-1^) of each forward and reverse primer and 2μl of undiluted DNA extract. The reaction conditions for PCR were an initial denaturation step at 95°C for 10 minutes followed by 35 cycles of 95°C for 30 seconds, 50°C for 30 seconds, and 72°C for 1 minute, a final extension at 72°C was performed for 10 minutes. Samples were cleaned using ExoSAP-IT Express (Applied Biosystems, California, USA) as manufacturer’s instructions. Successful PCR amplicons were Sanger sequenced with both primers at Eurofins Genomics (Ebersberg, Germany). The resulting sequences were quality trimmed and aligned. Only one primer provided good quality sequencing results (LCO1490). Both concatenated sequences provided full length (>90% query cover) excellent match quality (>97% identity) BLAST hits to multiple (>20) sequences corresponding to *A. senhousia* on the NCBI nt database corresponding with many independent studies (Genbank Accessions:AB498016.1, AY570034.1, HG005372.1, HQ891034.1).
